# Tuberculosis susceptibility and vaccine protection are independently controlled by host genotype

**DOI:** 10.1101/064253

**Authors:** Clare M. Smith, Megan K. Proulx, Andrew J. Olive, Dominick Laddy, Bibhuti B. Mishra, Caitlin Moss, Nuria Martinez Gutierrez, Michelle M. Bellerose, Palmira Barreira-Silva, Jia Yao Phuah, Richard E. Baker, Samuel M. Behar, Hardy Kornfeld, Thomas G. Evans, Gillian Beamer, Christopher M. Sassetti

**Author notes:** Address correspondence to: Christopher M. Sassetti. Department of Microbiology and Physiological Systems, University of Massachusetts Medical School, Worcester, MA 01605, USA.

## Abstract

The outcome of *Mycobacterium tuberculosis* (Mtb) infection and the immunological response to the Bacille Calmette Guerin (BCG) vaccine are highly variable in humans. Deciphering the relative importance of host genetics, environment, and vaccine preparation on BCG efficacy has proven difficult in natural populations. We developed a model system that captures the breadth of immunological responses observed in outbred individuals, which can be used to understand the contribution of host genetics to vaccine efficacy. This system employs a panel of highly-diverse inbred mouse strains, consisting of the founders and recombinant progeny of the “Collaborative Cross”. Unlike natural populations, the structure of this panel allows the serial evaluation of genetically-identical individuals and quantification of genotype-specific effects of interventions such as vaccination. When analyzed in the aggregate, our panel resembled natural populations in several important respects; the animals displayed a broad range of Mtb susceptibility, varied in their immunological response to infection, and were not durably protected by BCG vaccination. However, when analyzed at the genotype level, we found that these phenotypic differences were heritable. Mtb susceptibility varied between lines, from extreme sensitivity to progressive Mtb clearance. Similarly, only a minority of the genotypes was protected by vaccination. BCG efficacy was genetically separable from susceptibility, and the lack of efficacy in the aggregate analysis was driven by nonresponsive lines that mounted a qualitatively distinct response to infection. These observations support an important role for host genetic diversity in determining BCG efficacy, and provide a new resource to rationally develop more broadly efficacious vaccines.

**Importance:** Tuberculosis (TB) remains an urgent global health crisis, and the efficacy of the currently used TB vaccine, *M. bovis* BCG, is highly variable. The design of more broadly-efficacious vaccines depends on understanding the factors that limit the protection imparted by BCG. While these complex factors are difficult to disentangle in natural populations, we used a model population of mice to understand the role of host genetic composition to BCG efficacy. We found that the ability of BCG to protect an individual genotype was remarkably variable. BCG efficacy did not depend on the intrinsic susceptibility of the animal, but instead correlated with qualitative differences in the immune response to the pathogen. These studies suggest that host genetic polymorphism is a critical determinant of vaccine efficacy and provides a model system to develop interventions that will be useful in genetically diverse populations.

## Introduction

The outcome of an encounter with *Mycobacterium tuberculosis* (Mtb) is highly variable. Most individuals contain the infection and can remain asymptomatic for a lifetime. Only a fraction of infected individuals develop active disease; and even among these, the timing, location, and presentation of the pathology is remarkably diverse (1). The underlying basis of the variable outcome of Mtb infection is unknown and likely involves a complex interplay between environmental factors and genetic variation in both host and pathogen (2). Classic evidence for a role of host genetics driving disease outcome comes from twin studies showing a higher tuberculosis (TB) concordance rate in monozygotic compared to dizygotic twins (3, 4). More recently, linkage analyses defined rare Mendelian traits that cause extreme susceptibility to mycobacterial disease in children (5-9), and a variety of case-control (10, 11), linkage (12) or genome-wide association studies (13, 14) have implicated more common genetic variants in TB risk. The identification of these TB-associated polymorphisms provides valuable insight into the pathogenesis of this disease, as many of the identified genes function either in the establishment of a protective Th1-biased cell-mediated immune response (15), regulate disease-promoting inflammation (16, 17), or alter the pathogen’s intracellular environment (18). However, these known mechanisms explain only a small fraction of the variability observed in natural populations (2), suggesting an important role for interactions between these and other disease-modifying polymorphisms.

This diversity in TB susceptibility is mirrored in the variable efficacy of vaccination for this disease. The only TB vaccine that has been shown to protect humans is an attenuated strain of *M. bovis*, Bacille Calmette Geurin (BCG). This vaccine has been administered to more than three billion humans since it was developed in the 1920’s. In the subsequent decades, studies in different geographic regions and ethnic populations have produced widely variable estimates of BCG’s effect. In several populations, BCG efficacy is estimated to be greater than 75%. However, in regions where TB remains endemic, no significant protection from pulmonary TB in adults can be detected (19, 20). This variable efficacy could be due to differences in the vaccine strain or preparation, the environment, or the genetic background of the host. The roles of genetic variation in the vaccine strain (21-24) and previous exposure to environmental mycobacteria (25-27) have been investigated extensively. In contrast, the role of host genetic variation in BCG efficacy has been more difficult to quantify. It is clear that variations in many immune mediators are highly heritable (28-30). Many of these heritable variations affect mediators that are likely to be relevant to Mtb immunity, such as the number of central memory T cells or abundance of cytokines such as IL-12p40, GM-CSF, IFNa, and IL-6 (31). Indeed, numerous studies suggest that the immunological response to mycobacterial infection (32-35) and BCG vaccination (36, 37) is heritable. However, the relationship between these immunological markers and vaccine efficacy is unknown and very difficult to address in natural populations. Thus, while there is reason to suspect that BCG efficacy is influenced by genetic variation, it has proven difficult to dissociate these effects from other confounding variables. In particular, the effect of BCG is difficult to dissociate from the intrinsic TB susceptibility of each individual in a natural population.

In theory, animal models could be used to dissect the role of genetic diversity in vaccine protection. However, while the mouse model of TB has been very useful for understanding the mechanisms underlying Mendelian susceptibility to TB, this approach has proven less useful for understanding the complex trait genetics that that have been shown to underlie TB susceptibility in mice, with only 2 host loci so-far identified from forward-genetic approaches (38-40). A fundamental limitation of the classic inbred strains of *M. musculus domesticus* that are generally used to model TB is their genetic homogeneity, as 90-95% of these animals’ genomes are estimated to be functionally identical (41). As a result, these mouse strains mount qualitatively comparable immune responses to this infection and vary only modestly in their susceptibility to Mtb (42). In virtually every strain, Mtb grows logarithmically for 2-3 weeks, at which point bacterial growth is restricted by a strong Th1-biased CD4+ T cell response. Immunity depends largely on CD4+ T cell-derived IFNg, and allows the animal to survive for several months harboring a relatively constant burden of bacteria. BCG vaccination also produces a relatively homogenous effect in a variety of standard lab strains of mice (43-45), accelerating the initiation of adaptive immunity by several days, reducing the peak burden of bacteria by approximately 10-fold (46) and extending survival (47). It is often noted that the mouse model does not reproduce many aspects of TB disease seen in humans or non-human primates, including variable histopathology (48), progressive bacterial killing (49), and widely varying susceptibility ((e.g. 50). While these discrepancies are generally attributed to species difference, the lack of variation observed in mice could also reflect the low genetic diversity between classical inbred strains (51). Thus, the true range of TB-related traits that can be modeled in mice remains unclear.

Tractable model systems that incorporate relevant genetic diversity could be used to decipher the genetic determinants of both TB susceptibility and vaccine protection. Recently, a number of related model outbred mouse populations have been developed based on the same set of eight genetically diverse founder lines, which represent all three *M. musculus* subspecies. Outbred progeny of these strains, called the Diversity Outbred (DO) population represent similar genetic diversity as a human outbred population (52, 53) and display a remarkable heterogeneity in Mtb susceptibility (54). However, the completely outbred structure of this population imposes several limitations on the resource. Most notably, each genotype is represented by only a single animal. As a result, it is difficult to quantify the genotype-specific effect of an intervention, such as vaccination. To complement this resource, panel of recombinant inbred lines derived from the same founders was generated and called the “Collaborative Cross” (CC) (55, 56). The CC panel retains the genetic diversity of DO animals, but allows each genotype to be infinitely reproduced. We took advantage of the unique structure of the CC population to investigate the relationship between host genotype, TB susceptibility, and BCG efficacy. We found that the panel of parental and recombinant CC strains (hereafter named “diversity panel”) reproduced many aspects of outbred populations, and encompassed reproducible phenotypes that extend well beyond those observed in standard inbred strains. Using the diversity panel, we found that BCG efficacy is genetically dissociable from TB susceptibility and correlates with intrinsic immune biases in the strains. This new genetic resource identified an important role for host genetic diversity in vaccine efficacy and provides new approaches to understand the immunological basis of protection and to optimize vaccines for outbred populations.

## Results

### Genetically diverse mice display a broad range of phenotypes upon Mtb infection

In order to independently assess the contribution of host genotype to Mtb susceptibility and vaccine protection, we assembled a population of highly-diverse and reproducible inbred lines. The “diversity panel” contained the eight founder lines of the CC and DO populations, which include five relatively diverse classical inbred strains (A/J, C57BL/6J, 129S1/SvImJ, NOD/LtJ and NZO/H1LtJ) and 3 wild-derived inbred strains (CAST/EiJ, PWK/PhJ and WSB/EiJ), hereafter abbreviated to A/J, B6, 129, NOD, NZO, CAST, PWK and WSB. We also took advantage of additional recombinant CC lines, since the range observed for polygenic traits in this type of panel is often much greater in the genetically mosaic offspring than in the founders(57, 58). Recombinant lines were included in this panel based on their relative resistance or susceptibility to Mtb after intravenous challenge, which was determined in a related study.

To characterize the range of Mtb susceptibility represented in the diversity panel, we infected groups of each genotype with H37Rv via the aerosol route and monitored disease over time. When the population was analyzed in the aggregate (Fig 1A, B and C), we observed a range of susceptibility similar to that observed in completely outbred ‘DO’ animals derived from the same founders (54). Lung and spleen CFU burden varied over by range of 1000-10,000 fold, and some animals required euthanasia after only 4 weeks of infection. When the panel was assessed at the genotype level (Fig 1D, E and F), we found that these differences in susceptibility were reproducible within each genotype and were therefore highly heritable. Based on bacterial burden in the lung at 6 weeks post-infection, the genotypes could be ranked based on susceptibility (Fig 1D). The B6 strain, generally considered to be the most resistant of the classical inbred lines (59), showed intermediate susceptibility in our panel, maintaining a lung pathogen burden of approximately 10^6^ CFU throughout the infection. The CC recombinant line, CC001, proved to be the most resistant to infection by 6 weeks, while the wild derived line of *M. musculus domesticus*, WSB, was the most susceptible. The phenotypes of these outlier strains were notable. CC001 was the only line in the panel in which lung CFU burden decreased between weeks 3 and 6 post-infection (p=0.044 by *t*-test). This relatively modest decrease was the first indication that this strain progressively kills Mtb over time (as shown in the subsequent studies presented below, see Fig. S2). In contrast, the both WSB and CC042 strains were unable to control bacterial replication (p=0.0001 for WSB, p<0.0001 for CC042 compared to B6 via one-way-ANOVA with Tukey’s multiple comparison test), lost weight, (p=0.0039 for WSB, p=0.0004 for CC042 compared to B6 via one-way-ANOVA with Tukey’s multiple comparison test), and were moribund after only 4 weeks of infection. Thus, these genotypes were as susceptible as knockout mice lacking critical aspects of CD4+ T cell-driven immunity, such as NOS2 (60) or MHCII (61).

**Figure 1.**
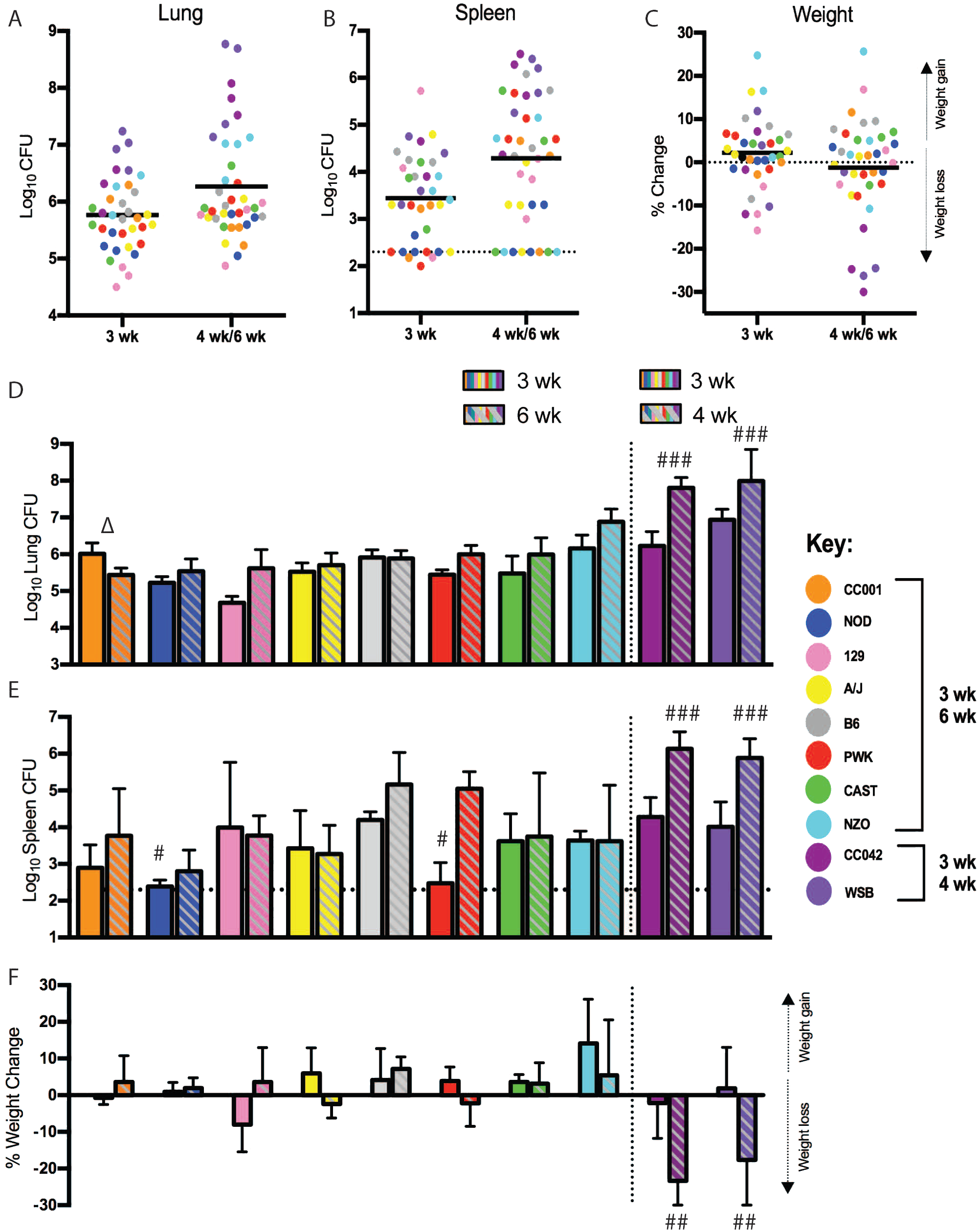
Mtb disease phenotypes in diverse mice. (A) Lung CFU (B) Spleen CFU and (C) weight change relative to initial weight for individual mice, colored by genotype at week 3 and 6 (for CC001, NOD, 129, A/J, B6, PWK, CAST and NZO) or week 3 and 4 (for WSB and CC042, which were moribund by wk 4). (D) average Lung CFU (E) average spleen CFU and (F) average weight change broken out by mouse genotype. All data is the average value of n=3 or 4 mice per strain at each time point, error bars are +/-SD. Statistical comparisons within mouse strains indicated by Δ (P<0.05) using student’s t-test, and # indicates statistical significance as compared to B6 via one-way-ANOVA with Tukey’s multiple comparison test (P<0.05#, P<0.01##, P<0.001###)

The ability to accurately measure multiple metrics of disease over time in reproducible lines, allowed the identification of traits that are genetically separable. For example, the rate at which bacteria disseminate from the lung to spleen was variable across the panel. Despite harboring similar numbers of Mtb in the lung at 3 weeks post-infection, NOD, and the wild-derived strain of *M. musculus musculus*, PWK, harbored significantly fewer CFU in their spleen than B6 (p = 0.020 for NOD vs B6 and p = 0.028 for PWK vs B6, by one-way ANOVA with Tukey’s multiple comparison test). In the spleens of PWK mice, Mtb burden increased between 3 and 6 weeks post-infection, indicating Mtb is able to replicate at this site. Thus, the early deficit in spleen CFU in this genotype could reflect delayed dissemination from the lung, which occurs during this period and has been previously shown to vary with host genotype (62). We conclude that the diversity panel encompasses a wide variation in TB susceptibility traits, and different aspects of disease may be controlled by distinct genetic polymorphisms.

### TB pathogenesis and immune response differ qualitatively between strains

The polygenic basis of susceptibility in the diversity panel suggested that distinct inflammatory and/or immunological pathways might underlie disease outcomes. Indeed, the lungs of infected mice displayed lesional heterogeneity between strains over time. At 3 and 6 weeks of Mtb infection, the lungs of B6 mice (the standard for mouse Mtb studies) contained typical lesions for this strain: multifocal coalescing histiocytic alveolar pneumonia with perivascular and peribronchiolar lymphocytic aggregates, no necrosis and diminishing neutrophils over time (Fig 2A-C). The lungs of several similarly resistant inbred strains (A/J and CAST) shared these characteristics. In contrast, after only 3 weeks of infection the susceptible WSB strain had already developed dense neutrophilic inflammatory infiltrates in small and large airways, which after one additional week of infection progressed to widespread necrosis of inflammatory cells and lung tissue, associated with morbidity. Interestingly, Mtb burden alone did not predict the extent of lung damage or the type of microscopic lesions in all cases. For example, the lungs of CC001 mice displayed early neutrophil recruitment and necrosis that was not apparent in other similarly resistant lines, and resolved by 6 weeks post-infection when the bacterial burden had decreased in this strain. Thus, the CC001 strain has some capacity to tolerate and resolve early necrosis and neutrophilic-mediated lung damage due to Mtb infection. Qualitative differences in the immune response were more apparent when cytokine levels were measured in the lung (Fig 2D-E). Immunity to Mtb in B6 mice depends on IFNg and TNF, which contribute to the effector function of CD4+ T cells (63, 64). Lung TNF levels were relatively consistent across the panel at 3 weeks post-infection, and were only elevated in the moribund WSB mice after 4 weeks. In contrast, IFNg levels in these mouse genotypes varied by nearly 20 fold at the early time point. The wild-derived lines PWK and CAST, and the recombinant CC042 expressed remarkably low levels of this cytokine that were at or below the limit of detection (LOD) of the assay (LOD = 17 pg/mL). Splenocytes from PWK and CAST both produced IFNg upon polyclonal stimulation *ex vivo* (Fig S1), indicating that these cells were capable of producing cytokine that was detectable in our assay. Thus, the lack of IFNg expression in the lung of PWK, CAST, and CC042 mice was not due to an inherent inability to express the cytokine. Instead, these lines appear to mount a distinct response to Mtb infection, and the relative resistance of the CAST and PWK lines indicates that these responses can be effective in the absence of high levels of IFNg.

**Figure 2.**
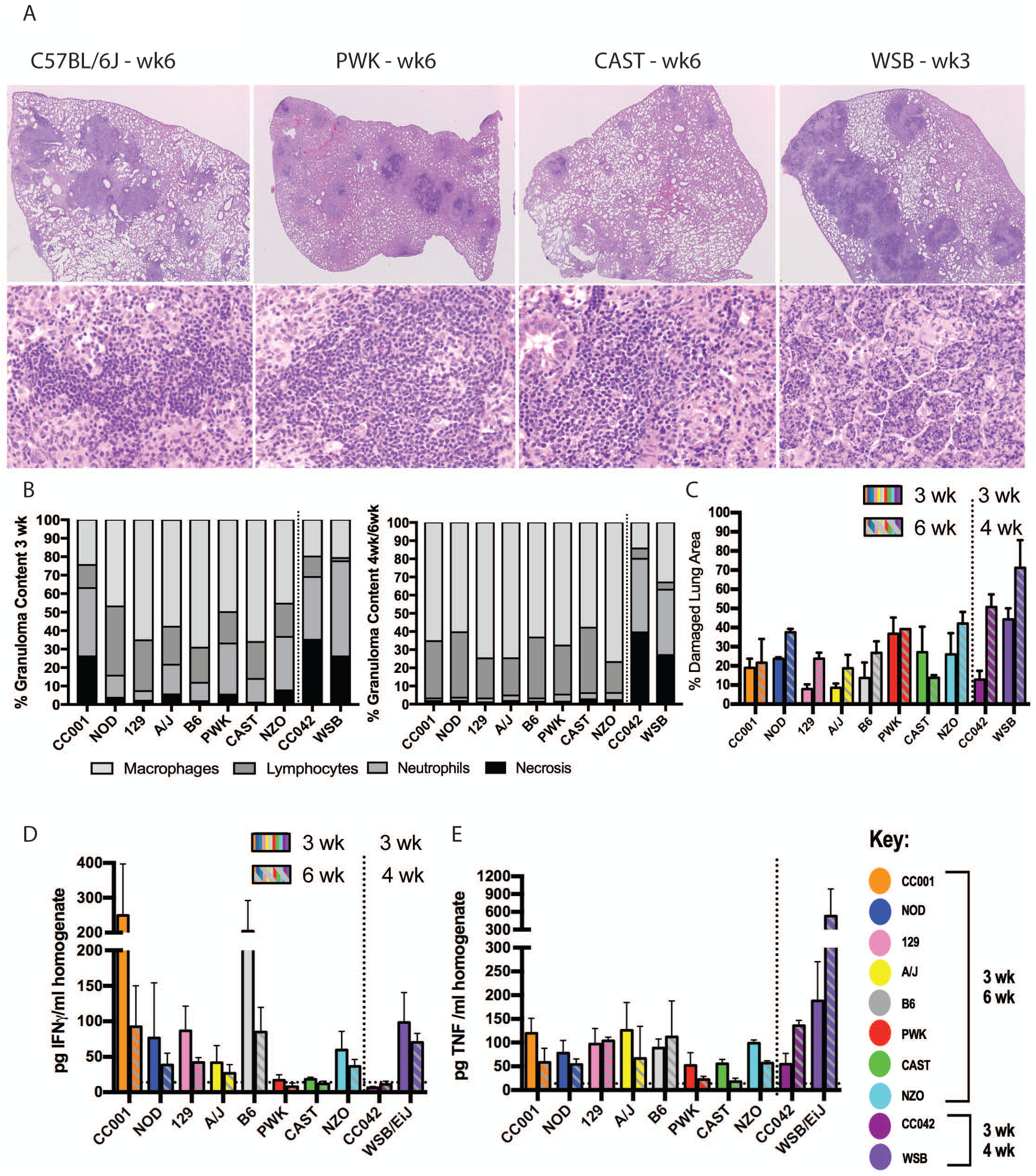
Distinct cytokine environment and histopathology in diverse mice. (A) Representative images from B6, PWK, CAST, and WSB mice at 2x and 20x magnification. (B) Proportional area of granuloma occupied by necrosis, neutrophils, lymphocytes and macrophages at 3 weeks post-infection (left) and 4wk or 6wk post-infection (right). Bar height is the average of 10 randomly selected granulomas for each of 3 to 4 mice per mouse strain. (C) Percent damaged lung area measured using ImageJ to trace lesion size relative to the whole lung section. Bar height is average of 3 or 4 mice per strain. Error bars are standard deviation. (D) IFNγ and (E) TNF cytokines measured in homogenates of infected lungs at 3 and 6 weeks (or 3 and 4 weeks for WSB and CC042) post infection. All data is average value of 3 or 4 mice per strain at each time point, assayed in technical duplicate, error bars are standard deviation. Limit of detection (LOD) calculated as 2-fold background (17 pg/ml).

### Host genotype determines BCG efficacy

To determine if genetic polymorphism could influence the degree of protection conferred by vaccination, we immunized a panel of diverse mouse strains with BCG by the subcutaneous route. The mice used for the vaccination study consisted of the same CC founder strains, the highly-resistant CC001 recombinant, and two additional recombinant lines representing susceptible (CC019) and resistant (CC002) phenotypes. Twelve weeks after vaccination, viable BCG was not detected in spleen homogenates of all genotypes (LOD = 20 CFU/spleen), indicating that all mice were able to control infection with this attenuated strain. Mice were then challenged with Mtb and the protection elicited by vaccination was determined at 4 and 14 weeks post-infection.

We initially analyzed this large panel as an aggregate population with 53-66 animals per group. As seen previously, the bacterial burden in lung and spleen of these animals varied widely, and vaccination did not alter this variation (Fig 3A-B). BCG vaccination reduced the mean Mtb burden in the lung or spleen by 1.7 to 4 fold at each time point, but even with this large group size the BCG-mediated reduction of bacterial burden was only statistically significant at 4 weeks post-infection. Thus, no durable protection from Mtb growth could be detected in the aggregated data, which mirrors the lack of protection conferred by BCG in many natural outbred populations.

**Figure 3.**
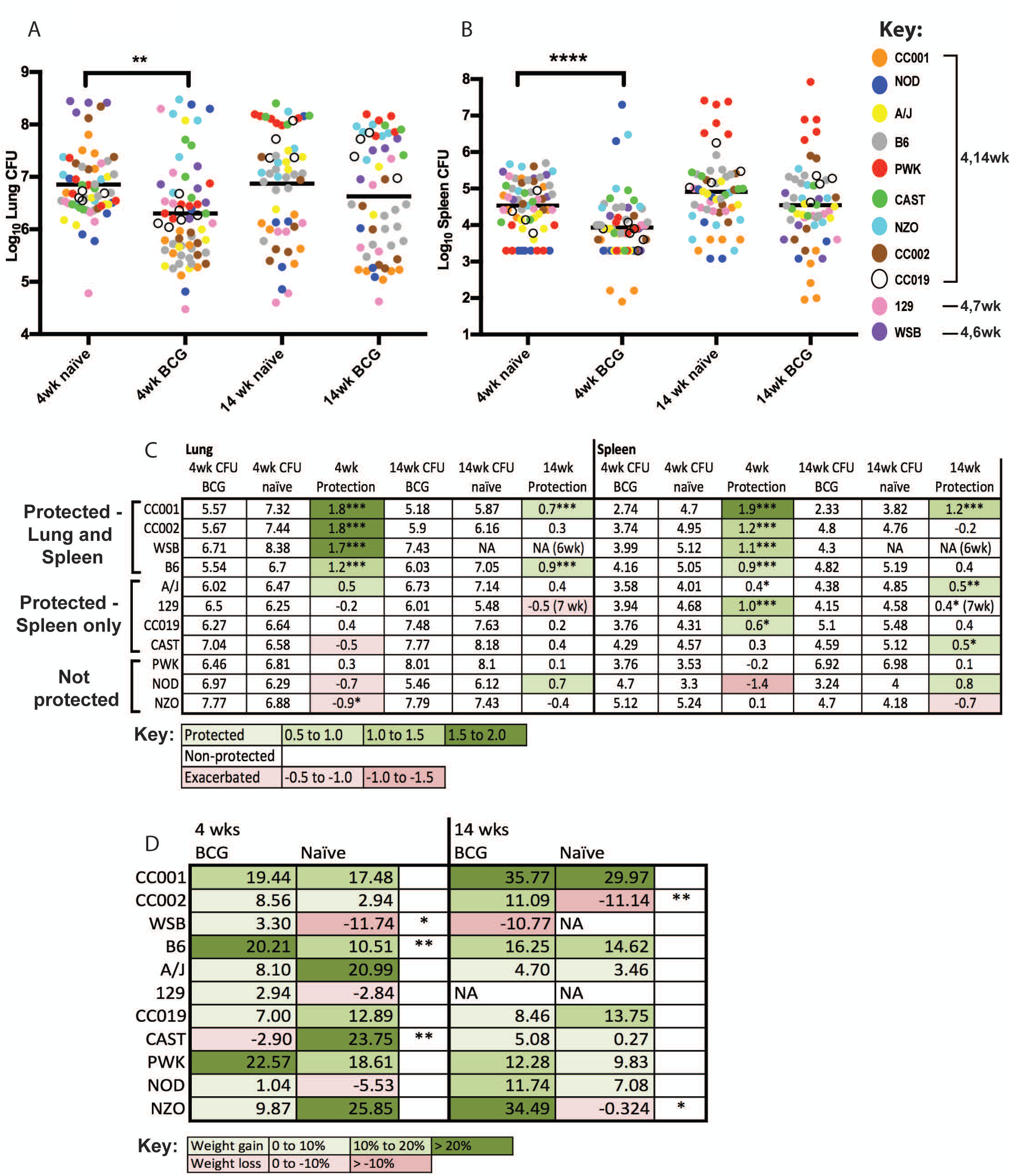
The effect of genetic background on BCG protection in diverse mice. (A) Lung CFU and (B) Spleen CFU in naïve and vaccinated mice for all genotypes. Each point represents the CFU for an individual mouse at the indicated time point. (C) Fold protection of each mouse strain was calculated from the average CFU of naïve compared to BCG-vaccinated groups at 4 and 14 weeks post-infection in the lung and spleen. Coloring indicates CFU reduction (green) or exacerbation (red) in the BCG vaccinated group. “Protection” is defined as a statistically significant 0.5 log_10_ reduction in the BGC vaccinated group. (D) Average % weight change of each mouse strain relative to initial body weight for BCG and naïve mice at 4 and 14 wks. The percentage of weight change was compared between naïve and BCG vaccination within each mouse strain (unpaired t-test). N=6 mice per strain per time point for naive or BCG-vaccinated. P<0.05*, P<0.01**, P<0.001***.

When the data were analyzed at the genotype level, it became clear that the variation in CFU burden was driven by genotype-specific effects on both Mtb susceptibility and vaccine efficacy (Fig. 3C-D). Using the reduction in Mtb burden as a metric of vaccine efficacy, we defined “protection” as a decrease in CFU of greater than 0.5 log_10_that reached statistical significance. Based on these criteria, only a subset of genotypes was protected by vaccination. Consistent with previous literature, BCG reduced the bacterial load in lungs and spleen of B6 mice by approximately ten-fold. This response was shared in three other lines, two recombinants (CC001 and CC002) and the wild-derived WSB line. In these genotypes, BCG exposure reduced Mtb load by 10-100 fold. Notably, these were the only genotypes in which vaccination protected the lung. BCG conferred protection in spleen but not lung of four additional lines (129, CC019, CAST and A/J). The remaining genotypes were not protected by vaccination in either organ at any time point. In these non-responding lines, BCG vaccination either had no effect on Mtb burden or was associated with an increase in mean CFU, which reached statistical significance for the NZO genotype (P<0.05, by t-test). In most cases, increased bacterial burden correlated with weight loss (Fig. 3D). In WSB, CC002, and B6 genotypes, BCG vaccination reduced bacterial burden and significantly reversed weight loss at one of the two time points. The effect of BCG on CFU and weight loss were discordant for the NZO and CC001 genotypes, reflecting unique phenotypes that will be discussed below.

#### TB susceptibility and BCG efficacy are genetically separable traits

The ability to serially evaluate the same host genotype allowed multiple traits to be independently measured in the vaccinated and unvaccinated state. To investigate potential mechanisms that determine vaccine efficacy, we searched for correlates of BCG-mediated protection among the traits measured during primary infection. These traits included lung cytokine measurements as a metric of adaptive immunity, CFU as a measure of antimicrobial capacity, and weight loss as a surrogate for systemic disease.

Upon correlating disease and immune metrics with the degree of BCG protection, we identified a number of clusters of covariant traits. A single cluster contained virtually all traits associated with BCG-mediated protection, including all metrics of antimicrobial efficacy and protection from weight loss at 4 weeks post-infection (Figure 4A, green box). The cluster did not contain any traits related to Mtb susceptibility in the unvaccinated state, and no specific correlation between Mtb susceptibility and BCG efficacy was evident (Fig 4B-C). Furthermore, BCG caused the largest reduction in CFU in CC001 and WSB, the most resistant and susceptible lines in the study, respectively (Fig. 4B, orange and purple). Thus, Mtb susceptibility and BCG efficacy were genetically separable.

**Figure 4.**
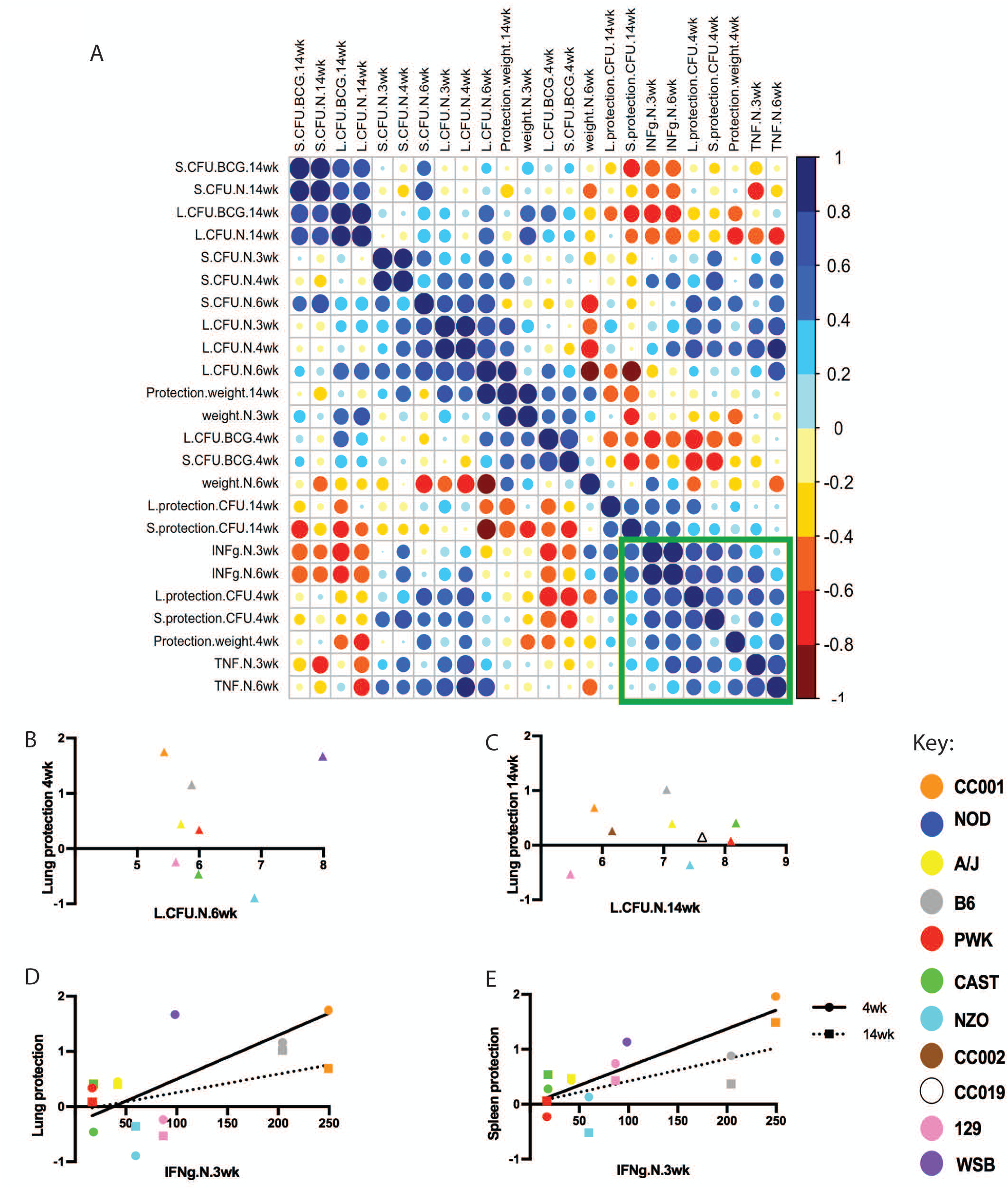
Phenotypic relationships between Mtb susceptibility and BCG efficacy. (A) Hierarchical clustering of correlations between metrics of susceptibility at week 3, 4, 6, and 14 post-infection and response to vaccination at week 4 and 14 post-infection. Blue indicates positive Pearson’s correlations and red indicates negative correlations. “Protection” indicates the relative reduction in CFU or weight loss in the BCG vaccinated group. “Weight” indicates % weight change relative to initial body weight. “N” and “BCG” indicate naïve or vaccinated groups, respectively. Green box indicates the trait cluster containing metrics of BCG-mediated protection. (B) and (C) Lack of correlation between TB susceptibility and BCG efficacy at early (B) or late (C) time points. (D) and (E) Positive correlation between IFNg production after Mtb infection and BCG efficacy in the lung (D) or spleen (E). Solid line is correlation of 4wk time point (circles), dotted line is correlation of 14wk time point (squares); Each data point in panels B-E is the average of 3 to 6 mice per genotype. Note that NOD mice were not displayed on the correlation plots due to the highly variable response of this strain (see Figure S2J).

In addition to all traits related to BCG protection, the BCG efficacy-related cluster of traits also included another metric of immune response that correlated strongly with the degree of BCG-mediated protection, the early production of IFNg by unvaccinated animals (Fig. 3D-E). The capacity to produce other T cell cytokines, such as TNF, was less predictive of BCG efficacy. Upon examination of individual genotypes, two of the wild-derived lines, CAST and PWK, are particularly notable for producing nearly undetectable levels of IFNg upon Mtb infection and receiving little or no benefit from BCG vaccination (Fig 4D-E, red and green).

### Long-term protection depends on a combination of traits

In this panel of genetically-diverse mice, we found that the extent of disease at later time points depended on the intrinsic susceptibility of the animal as much as the effect of vaccination. For example, despite the nearly 100-fold reduction in lung Mtb burden at 4 weeks post-infection, BCG only marginally extended the survival of the highly-susceptible WSB strain from 4 to 6 weeks, and ultimately did not protect this strain from disease. Conversely, even without vaccination, the resistant CC001 and CC002 strains had the remarkable ability to kill 90-95% of Mtb between 4 and 14 weeks postinfection (Fig 3 and Fig S2). As a result, unvaccinated animals of these genotypes harbored a similar bacterial burden at the later time point as the vaccinated B6 group. The relative resistance of this CC001 likely explains its ability to gain weight throughout the infection, regardless of BCG vaccination (Fig 3D).

We also found that mice could benefit from vaccination even if Mtb burden was not reduced. For example, even though BCG-vaccinated NZO mice harbored significantly more Mtb in their lungs than their unvaccinated counterparts at 4 weeks post-infection, vaccination still reversed Mtb-induced weight loss between 4 and 14 weeks (Fig 3D). The NZO mouse strain has been described to develop a polygenic form of type II diabetes at a similar age as the animals in our study (65). As chronic hyperglycemia can increase Mtb susceptibility in mice (66, 67), we hypothesized that an interaction between vaccination and diabetes initiation might underlie the paradoxical effect of BCG on Mtb burden and weight loss. Indeed, while 5 of 6 unvaccinated animals were found to have fasting blood glucose levels above 500 mg/dL at 14 weeks post-infection, 0 of 6 vaccinated animals were diabetic (Fig 5A-B). In the NZO genotype, diabetes strongly predicted weight loss but not CFU (Fig 5C-D). Our diversity mouse panel also included NOD mice that develop autoimmune type I diabetes (68). No significant effect of BCG on either Mtb burden or blood glucose concentration was evident in this strain, possibly due to the variable penetrance of diabetes (Fig S2J). Thus, the major protective effect of BCG in Mtb-infected NZO mice appears to be the prevention of type II diabetes. Taken together, our observations in the genetic diversity panel suggests that while the protective efficacy of BCG can be separated from TB susceptibility, the ultimate effect of vaccination on the outcome of infection is influenced by a variety of factors including intrinsic susceptibility to TB and complex interactions with co-morbidities such as diabetes.

**Figure 5.**
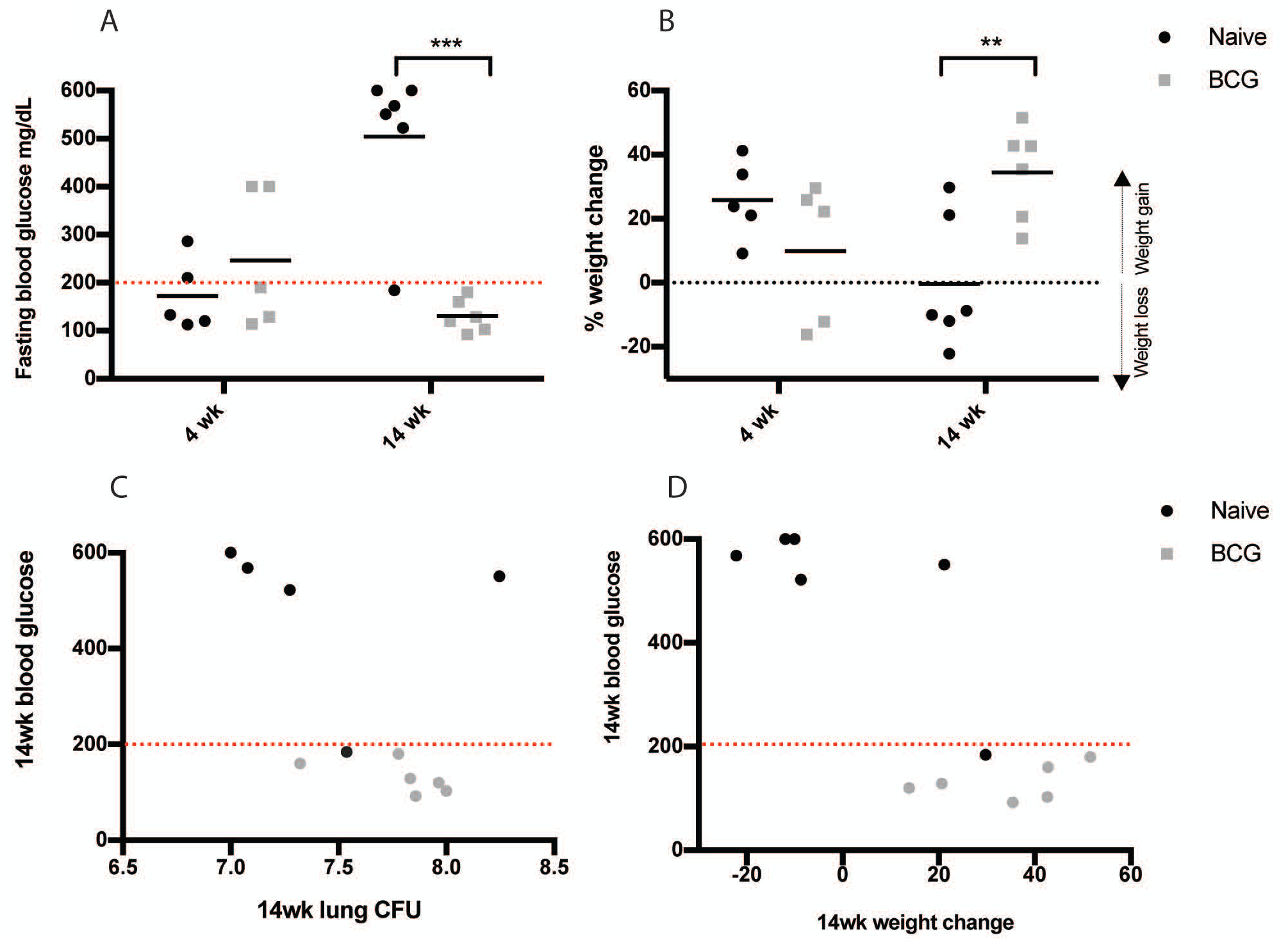
BCG protects Mtb-infected NZO mice from diabetes. (A) Blood glucose measurements obtained from naïve and BCG vaccinated NZO mice mice at 4 and 14 weeks post-infection. Mice were fasted for 6 hours prior to obtaining blood glucose measurements. Mice were considered diabetic if they had a blood glucose concentration of greater than 200mg/dL. (B) % weight change of naïve and BCG vaccinated mice relative to initial mouse weights at 4 and 14 weeks post-infection. (C) Correlation of blood glucose vs. lung CFU and (D) blood glucose vs. weight change at 14 weeks post-infection in naïve and BCG vaccinated mice. Each data point is an individual NZO mouse at each specified time point. Naïve and BCG vaccinated groups were compared by student’s t-test. P<0.05*, P<0.01**, P<0.001***

## Discussion

Predicting TB risk and rationally designing more effective interventions will ultimately require understanding the mechanisms that control the outcome of Mtb infection in a genetically diverse population. An ideal experimental model to dissect these mechanisms would encompass relevant genetic diversity and allow the serial evaluation of genetically identical individuals. To create such a population, we took advantage of the Collaborative Cross project, which identified genetically-diverse founder lines and generated recombinant inbred animals in which this diversity was reassorted to produce more extreme phenotypes. Our studies using this panel extend the phenotypic diversity described in the species and allowed the contribution of host genetics to TB susceptibility and BCG efficacy to be independently assessed.

The course of Mtb infection in the standard “mouse model” (B6 inbred strain) is often contrasted with the diversity of TB-related disease that is apparent in other species. For example, while Mtb can replicate continually at some sites in the lungs of cynomolgus macaques, Mtb is efficiently killed in most of the granulomas (49). These studies are consistent with observations from human autopsies, which similarly suggest that Mtb is eradicated from many granuloma and only replicates in a minority of sites (69). In contrast, with the exception of a few susceptible substrains (54, 70), Mtb infection of standard lab strains of *M. musculus domesticus* follows a very similar course; Mtb replication can generally be controlled, but the mice are unable to clear the pathogen or even to significantly reduce the lung bacterial burden during the persistent phase of infection when the Mtb burden remains stable. Our studies suggest that this homogeneity in TB pathogenesis reflects the genetic homogeneity of these strains, but not the phenotypic diversity of the species. Of course, the mouse lines used in this study may not capture all diversity previously described in mouse substrains. For example, the C3HeB/FeJ substrain forms caseating lung lesions upon Mtb infection (70), which closely resemble those of humans. Determining if any mouse lines in the CC panel recapitulate this phenotype will require additional infections studies in parallel with C3HeB/FeJ animals.

Phenotypes ranging from progressive killing of Mtb to extreme susceptibility were present in our panel, and appear to be based on different underlying genetic mechanisms. The ability to kill Mtb during the persistent phase of infection (between 4 and 14 weeks) was found only in recombinant animals and not the founder lines, suggesting that this is a multigenic trait that depends on the proper assortment of founder alleles. In contrast, several extreme phenotypes were evident in the more divergent wild-derived lines. The WSB line of *M. musculus domesticus* was extremely susceptible to Mtb. This line has also been recently reported to be modestly susceptible to influenza A virus infection (71). However, these animals are otherwise healthy and able to control the replication and dissemination of BCG, suggesting a fairly specific defect in immunity to Mtb. While the CAST line of *M. musculus castaneous*, and PWK line of *M. musculus musculus* appeared similarly susceptible to TB as many standard lines, their response to infection differed from other genotypes in several respects. The lungs of CAST mice displayed a paucity of inflammatory infiltrate, relative to other mice that harbored similar bacterial burdens, a trait also reported for this genotype during influenza virus infection (71). PWK animals were able to delay the dissemination of Mtb from the lung to the spleen. Most strikingly, both CAST and PWK produced barely detectable levels of IFNg during infection. While this cytokine is critical for Mtb immunity in B6 mice, these strains were still able to control bacterial replication. CD4+ T cells have been shown to control Mtb growth through IFNg-independent mechanisms in several settings (72, 73), and Th17 cells have been specifically implicated in protection (74). Together, these observations suggest that these highly divergent mouse lines might preferentially depend on non-Th1 biased CD4+ T cell responses for Mtb immunity.

The structure of our mouse population allowed us to determine that BCG efficacy is controlled independently of TB susceptibility in naïve animals. These observations suggest that even the most susceptible individuals can benefit from vaccination. However, the genetic basis of vaccine-conferred protection remains unclear. The animals in our panel encode diverse MHC haplotypes, raising the possibility that H2 polymorphism could influence the degree of protection (40). However, MHC haplotype alone is unlikely to explain the dramatic differences that we observed, since BCG produces a large and diverse array of potential antigens. In addition, two strains in our panel (129 and B6) shared the same H2 haplotype and were differentially protected, further indicating that additional mechanisms determine BCG efficacy. Similarly, while the relative ability to eradicate BCG could control efficacy, previous studies found that this vaccine protects animals of widely varying BCG susceptibilities, which were determined by Nramp1 (Slc11a1) polymorphism (44). Instead, the observed correlation between the propensity of an unvaccinated strain to produce IFNg after Mtb infection and the degree of BCG-induced protection suggests that intrinsic bias in the anti-mycobacterial immune response could influence vaccine efficacy. While this correlation was driven by both classical and wild-derived lines, the CAST and PWK were particularly strong outliers. These strains made very low levels of IFNg after infection and received virtually no benefit from BCG. Thus, it is possible that while BCG is a robust inducer of canonical anti-mycobacterial Th1 responses, it is unable to stimulate the immune response(s) that protect these highly-divergent strains.

The low and variable degree of protection elicited by BCG in natural populations could be due to either a general lack of efficacy or a differential effect in distinct individuals. It is important to understand the relative importance of these two factors, since entirely different approaches are necessary to overcome them. Our observations using a model population of genetically diverse mice indicate that genetic diversity in the host population could be a major factor limiting BCG efficacy. Based on these findings, it is not clear that optimizing a vaccine to protect a single standard lab strain of mouse will produce an intervention that is broadly efficacious in an outbred population, or even that a single vaccine is capable of protecting genetically diverse individuals. Instead, optimizing a vaccine or set of vaccines to protect non-responding strains may represent a more effective strategy. Furthermore, understanding the immunological biases that determine TB susceptibility vaccine efficacy would facilitate the identification of individuals that are genuinely at risk. The large panel of recombinant inbred Collaborative Cross lines that is currently available will allow the genetic dissection of these traits and should facilitate the development of more broadly effective vaccination strategies.

## Author contributions

CMS and CMS conceived and designed experiments. CMS, MKP, AJO, BBM, CM, MMB and JYP performed the experiments. CMS, MKP, GB, REB and CMS analyzed the data. NMG, HK, PBS, SMB, DL and TGE contributed reagents/materials/analysis tools. CMS and CMS wrote the paper.

## Acknowledgements

We thank Rustin R. Lovewell, Jarukit E. Long, and other lab members for technical assistance; Kadamba Papavinasasundaram and Jacqueline Schaeffer for helpful discussions and intellectual insight; and the UMASS Department of Animal Medicine for expert technical services.

## Materials and Methods

### Ethics statement

The animal studies were approved by the Institutional Animal Care and Use Committee at the University of Massachusetts Medical School (UMMS; Animal Welfare Assurance Number A3306-01), using the recommendations from the Guide for the Care and Use of Laboratory Animals of the National Institutes of Health and the Office of Laboratory Animal Welfare.

### Mice

C57BL/6J (stock #0664), A/J (stock #0646), 129SvImJ (stock #02448), NZO/HiLtJ (stock #02105), NOD/ShiLtJ (stock #01976), WSB/EiJ (stock #01145), PWK/PhJ (stock #3715) and CAST/EiJ (stock #0928) were purchased from the Jackson Laboratory (Bar Harbor, Maine, USA) and Collaborative Cross strains CC001, CC002, CC019, CC042 were purchased from the University of North Carolina (Chapel Hill, North Carolina, USA). Male mice were 8-12 weeks old at the start of all experiments. Mice were housed under specific pathogen-free conditions, and in accordance with the University of Massachusetts (UMASS) Medical School, IACUC guidelines.

### Vaccination

Mice were vaccinated with 10^5^ CFU BCG (BCG SSI strain; resuspended in 0.04% PBS-Tween80) via the sub cutaneous route (100uL per mouse). Mice were rested for 12 weeks; 3 mice per genotype were euthanized at 12 weeks post vaccination and spleens homogenized and plated to test for BCG persistence. 24 mice per genotype were subsequently infected with *M. tuberculosis*, n=12 vaccinated, n=12 naïve mice per genotype.

### Experimental infection and bacterial quantification

Infection with *M. tuberculosis* (*M. tuberculosis* H37Rv strain; PDIM positive) was performed via the aerosol route, with mice receiving 50-200 CFU. Bacteria were cultured in 7H9 medium containing 0.05% Tween 80 and OADC enrichment (Becton Dickinson). For infections, mycobacteria were suspended in phosphate-buffered saline (PBS)-Tween 80 (0.05%); clumps were dissociated by sonication, and inoculum delivered via the respiratory route using an aerosol generation device (GlasCol). To determine CFU, mice were anesthetized via inhalation with isoflurane (Piramal) and euthanized via cervical dislocation, the organs aseptically removed and individually homogenized, and viable bacteria enumerated by plating 10-fold serial dilutions of organ homogenates onto 7H10 agar plates. Plates were incubated at 37C, and *M. tuberculosis* colonies counted after 21 days.

### Cytokine measurement in tissue homogenates

Murine cytokine concentrations were measured from culture supernatants (prepared from uninfected splenocytes from n=3 mice per genotype; diluted to 1 million cells per well and stimulated by PMA/I or anti-CD3/CD28 for 72hrs) or cell free lung homogenates and quantified using commercial ELISA kits for IFNg and TNF (R&D systems, Duoset, Catalog #’s DY485, DY410, DY402). Final values are the average value from n=3 mice per genotype, with each individual mouse sample run in technical duplicate.

### Light microscopy

Lung lobes from 3 or 4 individual mice per time point were fixed with 10% neutral buffered formalin, processed, embedded in paraffin, sectioned at 5 µm, and stained with hematoxylin and eosin. Two serial sections were examined by a board certified veterinary pathologist (GB) for qualitative analyses including assessment of immune/inflammatory cell types; distribution and extent of lesions; presence and absence of necrosis; and extent and content of disrupted lung architecture. The extent of necrosis, neutrophils, macrophages and lymphocyte influx and was then estimated per strain per time point by blindly assessing 10 “granulomas” chosen at random, and reported as the proportional area occupied by necrosis, neutrophils, lymphocytes and macrophages.

### Statistical analysis

All data are represented as mean with SD. The statistical significance of differences between data groups was determined using unpaired two-tailed Student’s t-test, or genotypes compared to the standard B6 strain by one-way ANOVA with Tukey’s multiple comparison test using GraphPad Prism 7. Pearson’s correlation was used to determine correlation between measured traits and visualized using corrplot v0.77 (ordered by hclust) in R version 3.2.4

## Supporting information captions

**Supplementary Figure 1.**
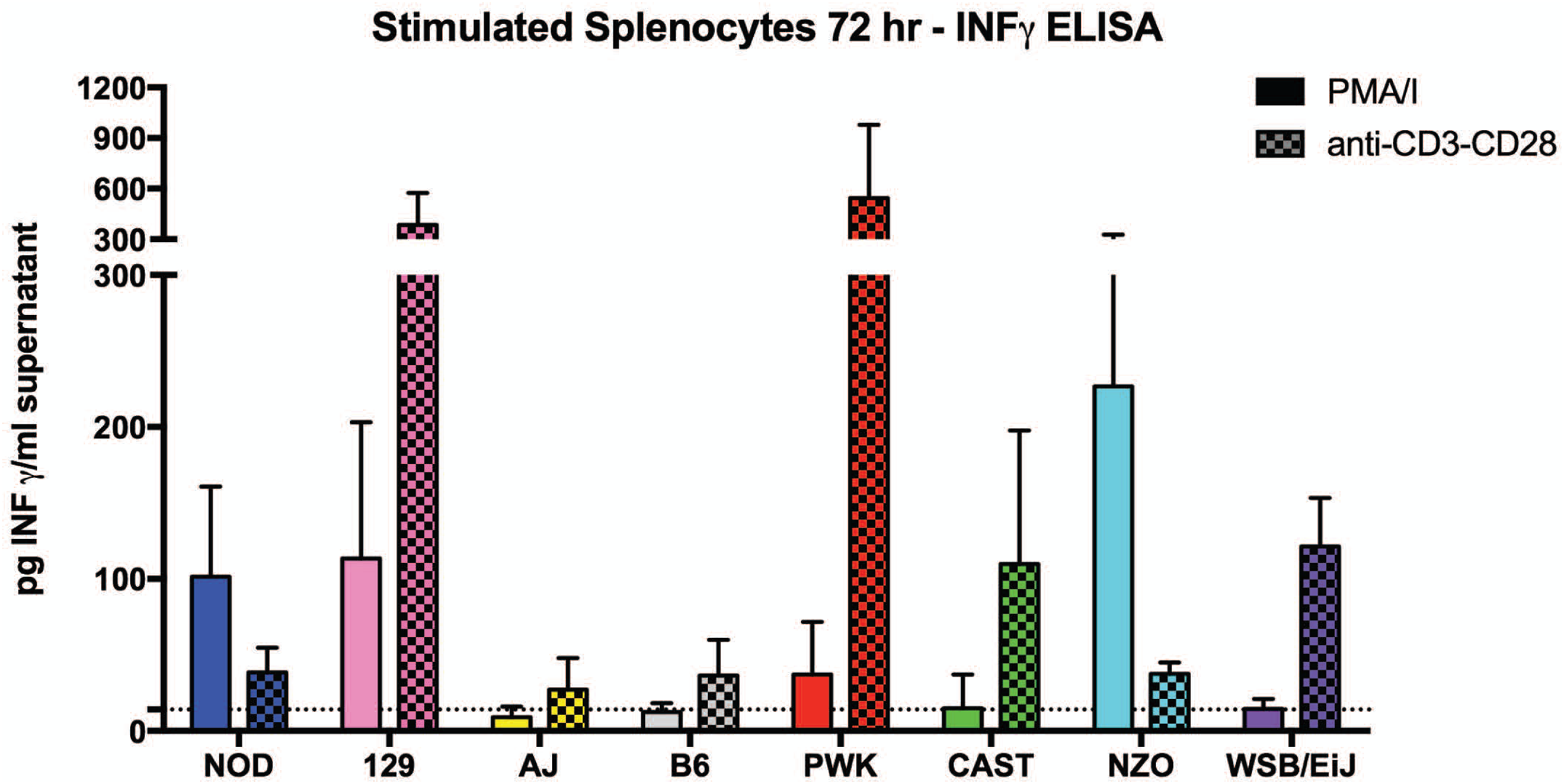
IFNg production *in vitro* from stimulated splenocytes. Spleens were collected from uninfected parental strains, homogenized, splenocytes counted, plated, stimulated with either PMA/I or anti-CD3/CD28, and incubated for 72hrs at 37°C before supernatants were collected and IFNg was measured by ELISA. Each bar is the average of 3 mice, with ELISAs performed in technical duplicate. Error bars are standard deviation.

**Supplementary Figure 2.**
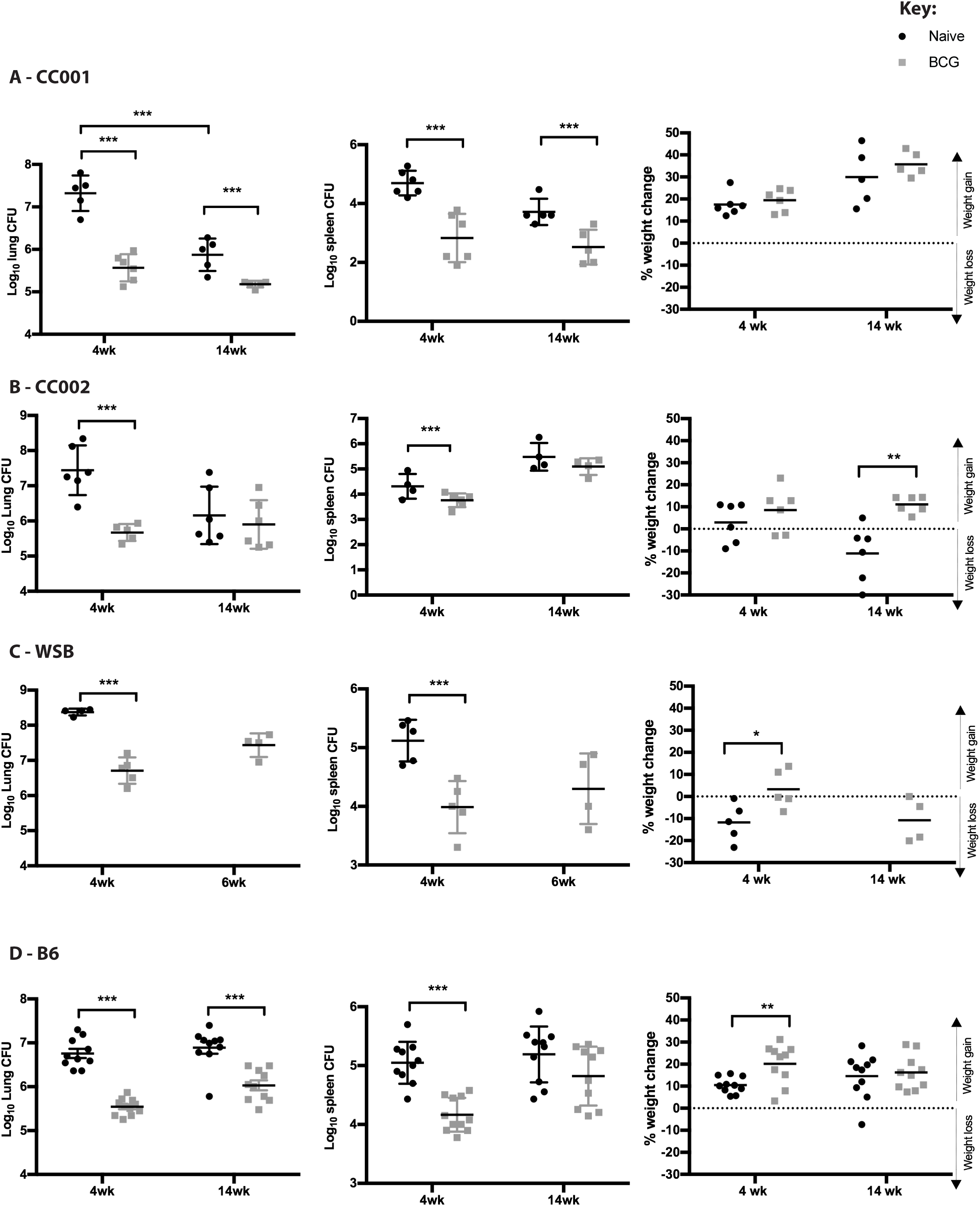
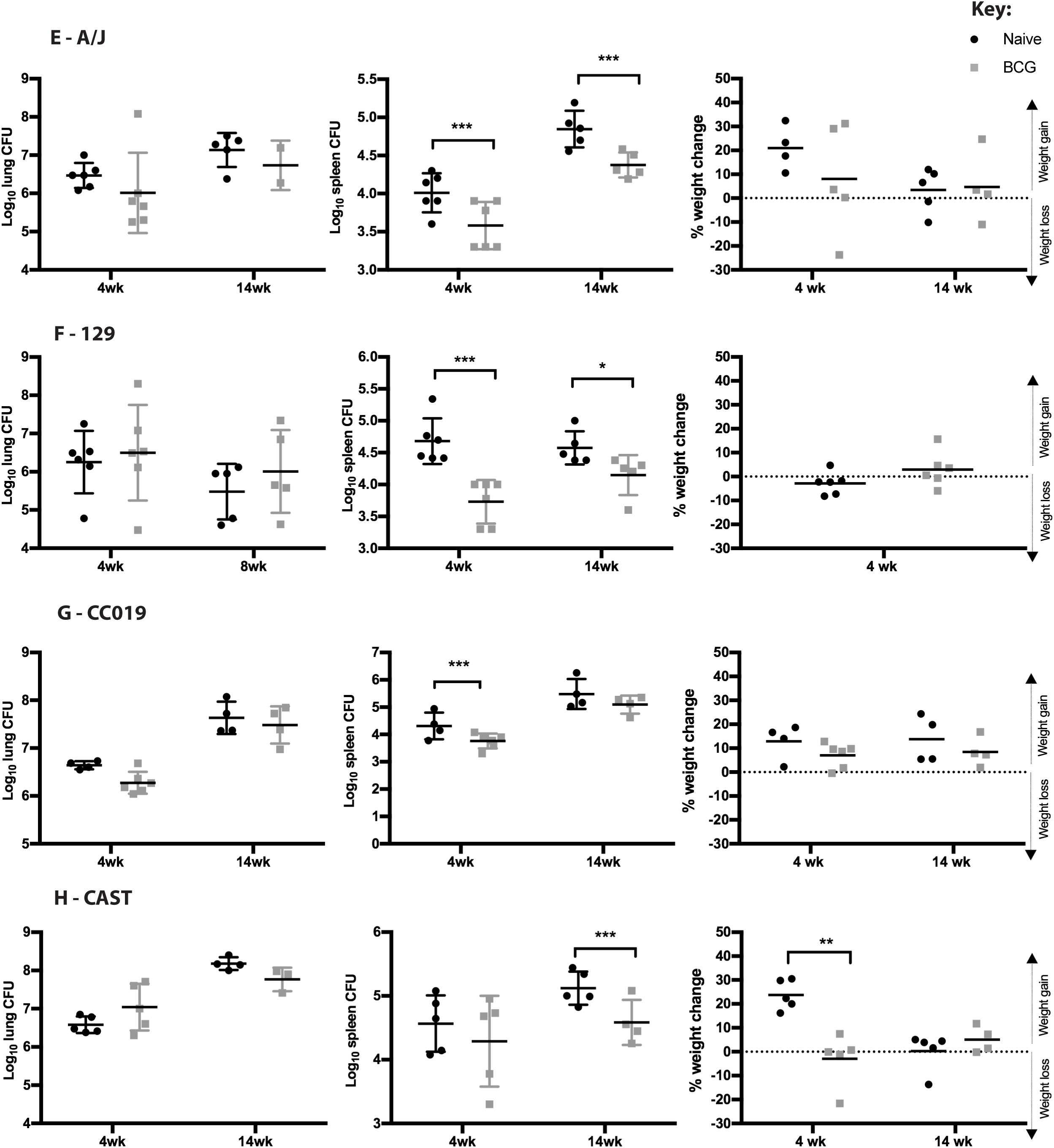
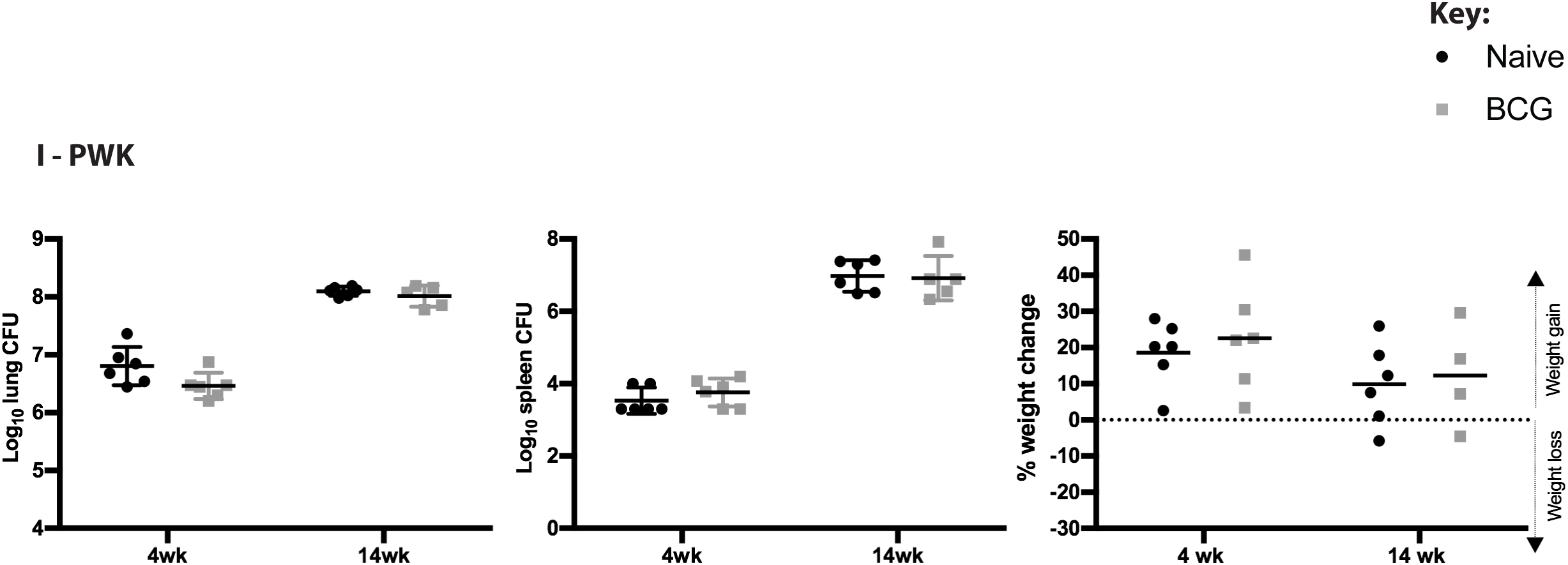
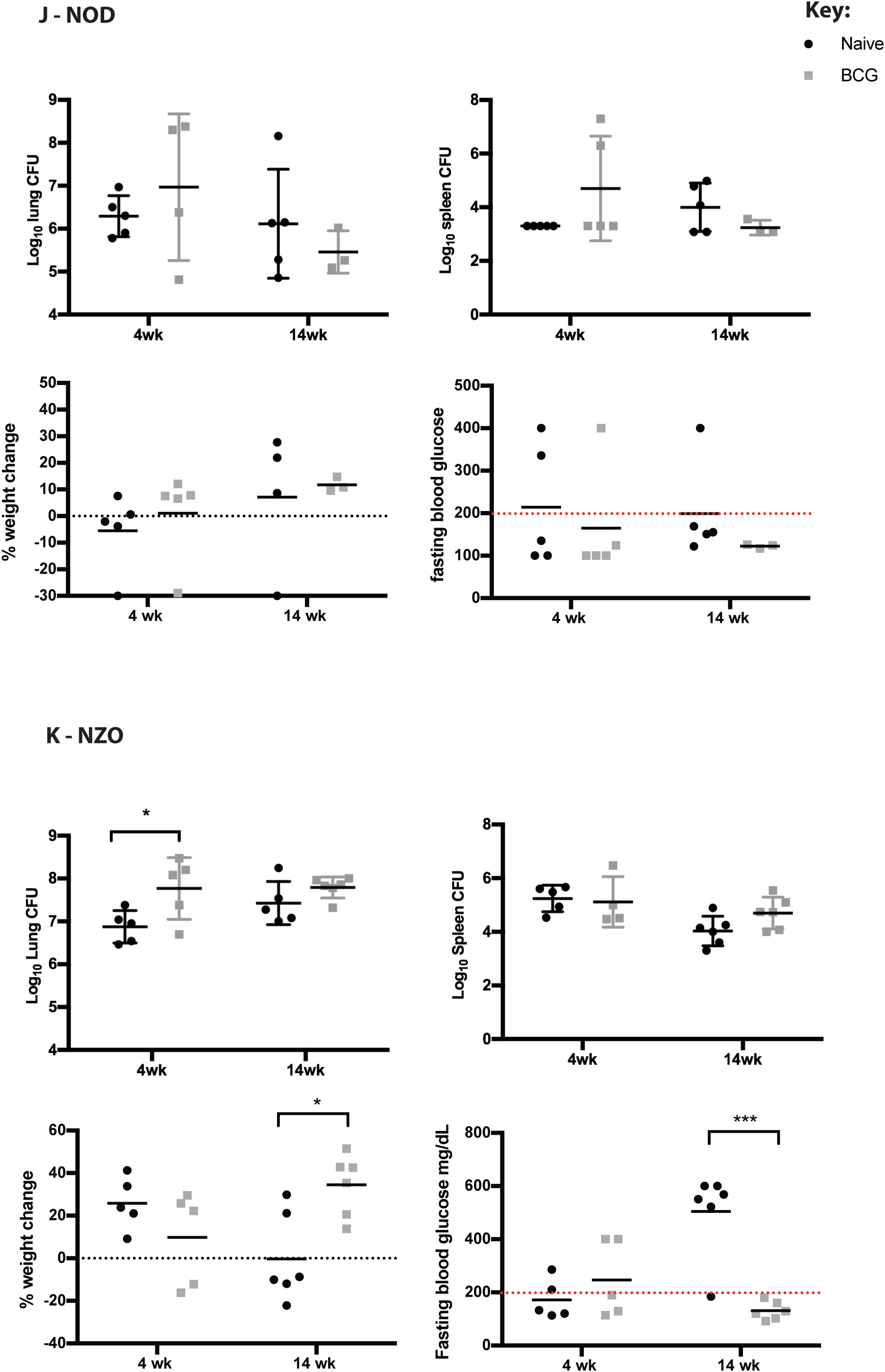
Lung CFU, Spleen CFU and weight change effects from BCG-vaccination by individual mouse genotype. Mice were vaccinated with BCG (n=12 per genotype) or left naïve (n=12 per genotype) and rested for 12 weeks prior to being infected with *M. tuberculosis*. Mice were weighed and euthanized at 4 and 14 weeks (6 mice per genotype, per treatment condition at each time point) and lung and spleen CFU were enumerated. Protection was calculated by comparing average CFU between naïve and BCG vaccinated groups. Statistical significance was determined by unpaired t-test (P<0.05*, P<0.01**, P<0.001***).

